# Spindly is Required for Rapid Migration of Human Cells

**DOI:** 10.1101/248393

**Authors:** Claudia Conte, Michelle A. Baird, Michael W. Davidson, Eric R. Griffis

**Affiliations:** University of Dundee, College of Life Sciences, Centre for Gene Regulation and Expression, Dundee, DD1 5EH, UK; Department of Biological Science, National High Magnetic Field Laboratory, Florida State University, Tallahassee, Florida, 32306, USA

**Author notes:** Current Address: Centre for Genomic Regulation (CRG), The Barcelona Institute of Science and Technology, 08003 Barcelona, Spain. Current Address: Nikon Instruments Inc, 1300 Walt Whitman Road, Melville, NY 11747, USA.

## Abstract

Dynein is the sole minus-end directed microtubule motor found in animals. It has roles in cell division, membrane trafficking, and cell migration. Together with dynactin, dynein regulates centrosomal orientation to establish and maintain cell polarity, controls focal adhesion turnover and anchors microtubules at the leading edge. In higher eukaryotes, dynein/dynactin requires additional components such as Bicaudal D to form an active motor complex and for regulating its cellular localization. Spindly is a protein that targets dynein/dynactin to kinetochores in mitosis and can activate its motility *in vitro*. However, no role for Spindly in interphase dynein/dynactin function has been found. We show that Spindly binds to the cell cortex and microtubule tips and colocalizes with dynein/dynactin at the leading edge of migrating U2OS cells and primary fibroblasts. U2OS cells that lack Spindly migrated slower in 2D than control cells, although centrosome polarization appeared to happen properly in the absence of Spindly. Re-expression of Spindly rescues migration, but the expression of a mutant, which is defective for dynactin binding, failed to rescue this defect. Taken together, these data demonstrate that Spindly plays an important role in mediating a subset of dynein/dynactin’s function in cell migration.

## Introduction

Cell migration is required for development and homeostasis in almost all multi-cellular organisms. The activation of this process requires specific stimuli from growth factors, chemokines or extracellular matrix molecules, which activate specific receptors and signalling cascades (Ridley, 2011). To migrate in 2-dimensions, a cell must protrude its plasma membrane, anchor these protrusions to the underlying substrate, and then use these connections to pull the cell body forward while constricting the rear of the cell and disassembling and releasing old connections (Krause and Gautreau, 2014). This process therefore requires the careful coordination between adhesive complexes, cytoskeletal filament systems with their attendant motor proteins, the secretory/membrane transport machinery and the regulatory molecules that control the activities of these disparate networks (Schmoranzer et al., 2003, Kaverina and Straube, 2011, Huber et al., 2015).

Actin microfilaments and non-muscle myosin II provide the majority of forces that drive migration. At the leading edge, small GTPases control the nucleation of actin filaments through the Arp2/3 complex, which produces branched filaments, and formin/Spire/JMY proteins, which build unbranched filaments (Ridley, 2011, Campellone and Welch, 2010, Firat-Karalar and Welch, 2011). The polymerization of actin at the leading edge of migrating cells generates the force to drive the extension of the cell membrane. Within the cell, bundled actin filaments, attached to focal adhesions, provide cables for the generation of traction forces that propel the cell body forward; at the rear of the cell, myosin contraction of actin filaments leads to the retraction of the cell body and release of focal adhesions (Ridley, 2011, Parsons et al., 2010, Aratyn-Schaus et al., 2011).

Although they do not themselves generate forces, microtubules are essential in many cell types for polarization and for regulating the speed of migration, and there are many points of feedback where microtubules, focal adhesions, and the actin network influence each other (Vasiliev et al., 1970, Kaverina and Straube, 2011, Stehbens and Wittmann, 2012, Akhshi et al., 2014). After an initial migration cue is received in a fibroblast, the microtubule organizing center (MTOC) orients itself between the nucleus and a cell’s leading edge and projects dynamic microtubules towards the lamellipodium (Magdalena et al., 2003, Gomes et al., 2005, Gotlieb et al., 1981, Gotlieb et al., 1983). These microtubules provide the tracks upon which membrane vesicles and locally translated mRNAs travel and deliver GTPase regulating proteins that activate Rac and Rho to stimulate focal adhesion internalization, actin polymerization or cell contraction (Rogers et al., 2004, Krendel et al., 2002, Rooney et al., 2010, Waterman-Storer et al., 1999, Montenegro-Venegas et al., 2010, Yadav et al., 2009, Mingle et al., 2005). There are also direct interactions between the actin cytoskeleton and microtubules through proteins such as the spectraplakin ACF7/MACF1 that can link microtubules to focal adhesions, the formin mDia1 that nucleates actin filaments and also stabilizes microtubules, and IQGAP1, which binds to several microtubule plus-tip proteins and can regulate actin and myosin activities (Palazzo et al., 2001a, Bernier et al., 2000, Karakesisoglou et al., 2000, Brandt and Grosse, 2007). Finally, motors of the kinesin superfamily are able to regulate microtubule dynamics, network architecture, and cargo transport and therefore many of them have roles in cell migration (for review see (Bachmann and Straube, 2015)).

As the only processive minus-end-directed microtubule motor, the dynein/dynactin supercomplex also has well established roles in cell migration. Dynein is targeted to growing microtubule plus-ends via the p150 subunit of dynactin, which is itself recruited by proteins, such as EB1 and CLIP-170, that bind to the plus-ends of microtubules and regulate their dynamics (Folker et al., 2005, Duellberg et al., 2014, Valetti et al., 1999, Vaughan et al., 1999, Vaughan et al., 2002). The dynein/dynactin complex was also described to be involved in cytoskeleton reorganisation upon wounding and in directed cell movement (Palazzo et al., 2001b, Faulkner et al., 2000, Smith et al., 2000). In addition to the dynein and dynactin complexes, several accessory factors, such as Lis1 and Ndel1, are important for the activity of the motor in many contexts, including cell migration (Cianfrocco et al., 2015, Reiner et al., 1993). Recent research has shown that vertebrate dynein and dynactin do not form a processive motor complex without activating factors such as Bicaudal-D (BicD) or Hook3. These activating factors drive dynein/dynactin supercomplex formation and allow it to move on microtubules (McKenney et al., 2014, Schlager et al., 2014, Urnavicius et al., 2015). Spindly has been shown to be one such activating factors (McKenney et al., 2014).

Spindly was identified through two RNAi screens in *Drosophila melanogaster* S2 cells in which mitotic and interphase phenotypes were analysed. In interphase cells, Spindly depletion generated alterations in cytoskeletal architecture with spiky and elongated microtubule-rich projections in contrast to the normal smooth rounded S2 cells. Moreover, GFP-Spindly was shown to track on the plus-ends of interphase microtubules, where it colocalized with the canonical plus-end binding protein EB1 (Griffis et al., 2007).

After the initial study in 2007, all of the subsequent publications on Spindly have been focused on describing its role during mitosis in human cells and worms (Gassmann et al., 2008, Gassmann et al., 2010, Holland et al., 2015, Yamamoto et al., 2008, Barisic et al., 2010, Cheerambathur et al., 2013, Chan et al., 2009, Moudgil et al., 2015); thus it was unclear whether Spindly in other organisms plays any functions in interphase cells.

In this study, we identified a direct role of human Spindly in wound healing and cell movement. Although predominantly a nuclear protein, Spindly localizes at the leading edge and focal adhesions in migratory cells. Cells lacking Spindly are slow to migrate in a scratch-wound assay, a defect that can be rescued by the reintroduction of the wild-type protein but not by the expression of a mutant that fails to bind to dynactin. Therefore, we can conclude that Spindly’s role in cell migration is likely due to its function in regulating dynein/dynactin activity, similarly to its established role in mitosis. These results delineate for the first time an interphase role for Spindly and confirm that this protein is a key adaptor for the dynein/dynactin motor complex in multiple cellular processes and in different cell cycle phases.

## Results and Discussion

### Localisation of human Spindly in fixed non-mitotic cells

To date, there has been very little data on human Spindly in non-mitotic cells, and so we began by assessing its localization. When we used an affinity-purified antibody raised against the full-length recombinant protein to stain U2OS cells that were grown in a monolayer and then scratched to induce cell migration, we noticed that, in addition to the expected nuclear staining, there was also a cytoplasmic pool of protein (Figure 1A upper). We confirmed the specificity of this staining by observing that siRNA depletion of Spindly eliminated the staining (Figure 1A lower and 1B). Fractionation of cells into nuclear and cytoplasmic fractions followed by western blotting demonstrated the presence of Spindly in both compartments (Figure 1C and Supplemental Figure S1).

**Figure 1.**
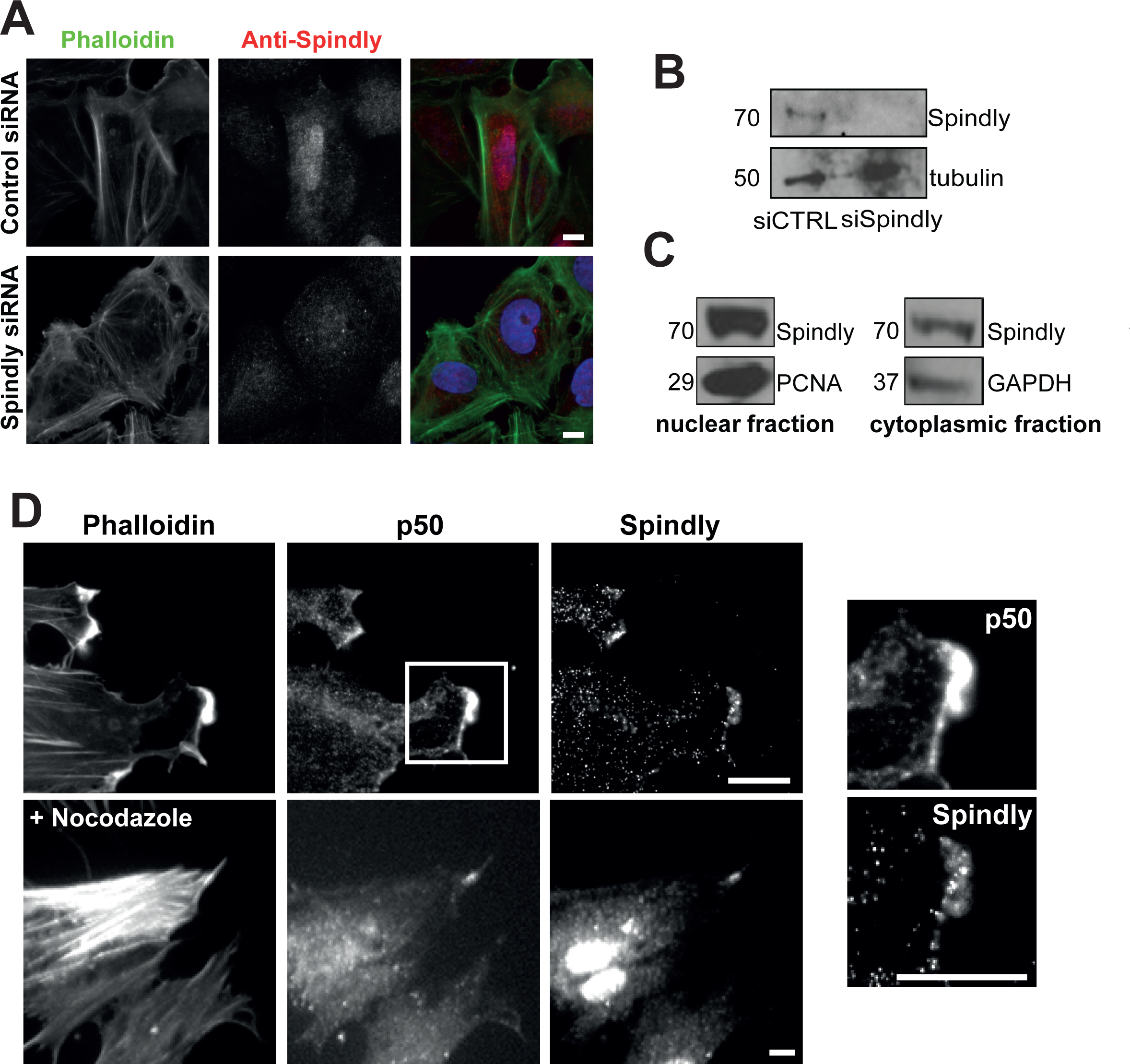
Spindly localizes to the leading edge of fixed migrating cells. A) Confluent U2OS cells were treated with control or Spindly-specific siRNAs and then cells were fixed and stained to visualize nuclei (DAPI), filamentous actin (phalloidin) and Spindly. B) An immunoblot of cell lysates show that Spindly was efficiently depleted by the siRNAs. C) U2OS cells were lysed and the cytoplasmic and nuclear fractions were separated. Co-fractionation with PCNA confirms Spindly presence in the nucleus and co-fractionation with GAPDH confirms the presence of Spindly in the cytoplasm. D) Foreskin fibroblasts were cultured to confluency, and then the monolayer was scratched to promote cell migration. Four hours after scratch-wounding, cells were fixed and stained to visualize filamentous actin (phalloidin), p50 Dynamitin, and Spindly. Images on the left show a magnification of the box shown in the upper image. Nocodazole treatment did not abolish the colocalization of p50 and Spindly. Scale bar = 10 μm.

To examine Spindly’s localization in a more migratory cell type and to determine if it localizes with any components of the dynein/dynactin complex, we fixed and stained primary human fibroblasts to visualize filamentous actin, the p50-Dynamitin subunit of dynactin and Spindly (Figure 1D). We clearly observed that Spindly and p50 co-localized at the leading edge of these cells (Figure 1D, lower panels). This colocalization was abolished by the application of latrunculin D (Supplemental Figure 2), but remained in cells treated with nocodazole to depolymerize microtubules (Figure 1E), suggesting that the proteins were associating with an actin-based structure.

### Live-cell imaging reveals that Spindly localizes to microtubule tips and mature focal adhesions

To further explore Spindly’s localization in interphase we asked whether Spindly could be seen associating with the basal cell cortex and/or cytoskeletal elements. We therefore imaged U2OS cells stably and inducibly expressing low levels of GFP-Spindly using total internal reflection fluorescence (TIRF) microscopy. TIRF allowed us to strictly visualize the localization of Spindly on or near the cell cortex, without interference from the nuclear signal, which is dominant in wide-field microscopy. In TIRF, we observed that there was a consistently bright fluorescent signal at the basal cortex. Additionally, we observed multiple populations of GFP-Spindly foci on the cortex. Some appeared to be moving diffusively, while others were more stable over multiple frames and appeared to be on cytoskeletal structures and in cytoplasmic projections (Figure 2A and Movie S1). In order to better understand how Spindly localizes in migrating cells, we visualized Spindly in U2OS cells that were grown to confluency and then scratch-wounded to induce cell migration. We observed that Spindly associated with the basal surface of the expanding plasma membrane and in rapidly moving foci (Movie S2).

**Figure 2.**
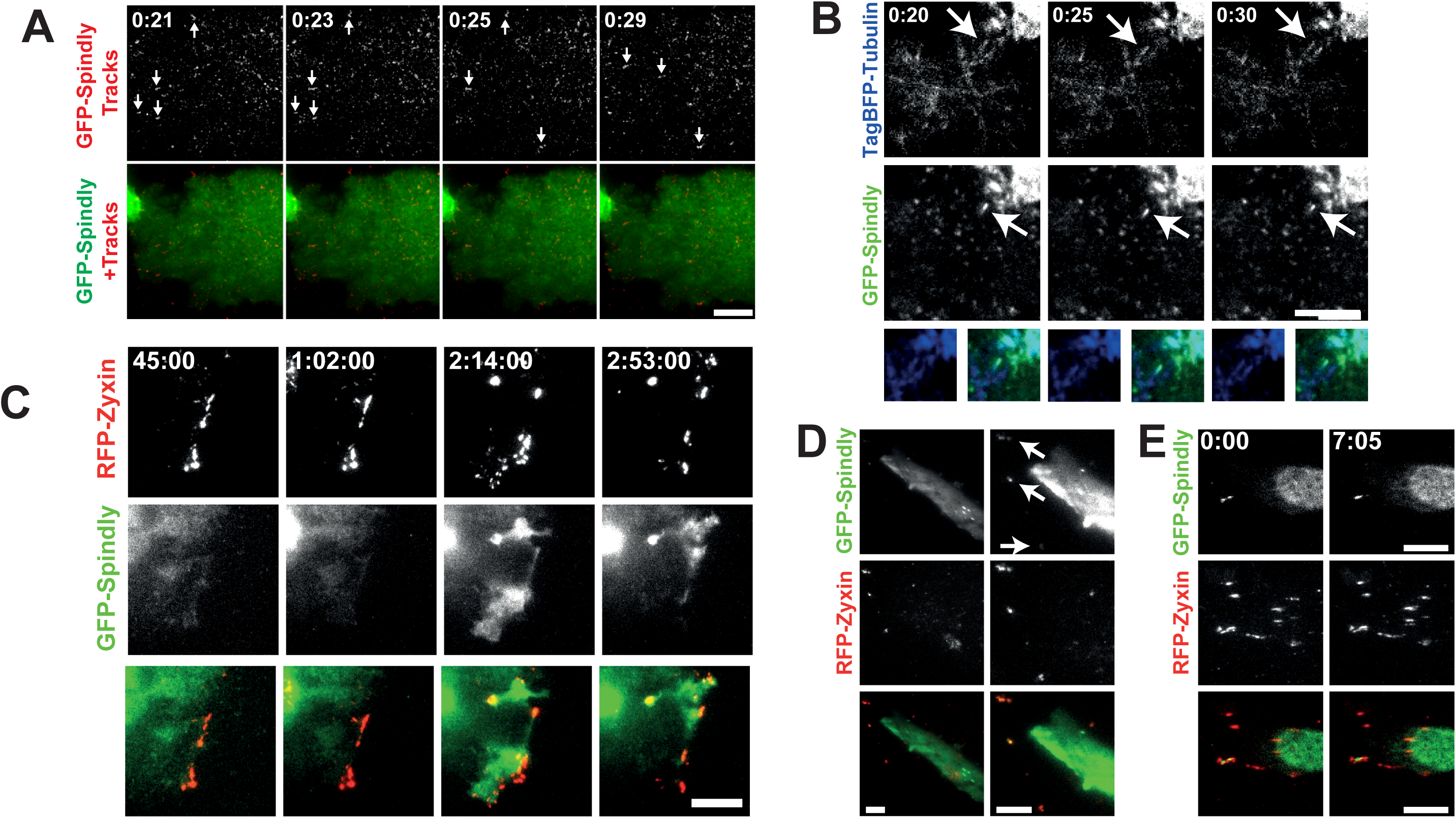
Spindly can be seen moving on the basal cell cortex and associated with microtubule plus-ends. A) U2OS cells stably expressing low levels of GFP-Spindly were imaged with total internal reflection fluorescence microscopy (TIRF). To better visualize the movement of individual particles across multiple images, a Gaussian filter was applied to denoise the images, and then a temporal filter was used to find foci that were moving across multiple images that were compiled into trails. Arrows show moving GFP-Spindly. The lower image shows the particle trails (red) overlaid on top of the still GFP-spindly image. Time is minutes:seconds. Scale bar = 10μm. B) The same U2OS cells as in A were transfected with TagBFP2-Tubulin and imaged in TIRF. The arrow shows a TagBFP2-labeled microtubule that shrinks and grows over the course of imaging. Time is minutes:seconds and scale bar = 10μm. C) The same U2OS cells were transfected with RFP-Zyxin, grown to confluency and then induced to migrate. GFP-Spindly only began to co-localize at the leading edge at 45 minutes post scratch-wounding. Time is shown in hours:minutes:seconds and the scale bar = 10μm. D) Fibroblasts were transiently transfected with GFP-Spindly and RFP-Zyxin and then imaged in TIRF. Arrows show focal adhesions where Spindly and Zyxin colocalize, which are more obvious in the first frame where the brightness was enhanced relative to the other time points. The arrowhead shows filaments of GFP-Spindly. E) In another fibroblast, a focus of GFP-Spindly was observed (arrowhead) that was dynamically associated with focal adhesions. Time is shown in minutes: seconds and the scale bar = 10μm.

To determine if the foci of Spindly were associated with microtubule plus-ends, we transfected our GFP-Spindly expressing U2OS cell line with TagBFP2-Tubulin and imaged these cells with TIRF. A small fraction of the observed particles of GFP-Spindly were clearly associated with microtubule plus-ends. The example shown in Figure 2B and Movie S3 contains a Spindly focus that remains on a microtubule that retracts and then grows over 10 seconds.

To further analyze the leading edge localisation of Spindly and its dynamics, we co-transfected our U2OS cells with RFP-Zyxin to visualize focal adhesions. Upon scratch-wounding, we followed the cells using TIRF microscopy. Figure 2C shows the recruitment of Spindly to the leading edge in these moving cells. We observed that it enriched with Zyxin, but only significantly after migration and Zyxin redistribution had begun. This indicates that Spindly could potentially require focal adhesion maturation to be recruited or be involved in the later stages of focal adhesion maturation or turnover (Nagano et al., 2012). Furthermore, in fixed cells we also observed the colocalization of endogenous Spindly and RFP-Zyxin at focal adhesions at the cell periphery (Supplemental Figure 3). When we cotransfected GFP-Spindly and RFP-Zyxin into resting fibroblasts, we also observed that some GFP-Spindly colocalized with RFP-Zyxin at focal adhesions (Figure 2D). However, the loss of Spindly did not seem to affect the size, number, or distribution of focal adhesions in U2OS cells (data not shown).

**Figure 3.**
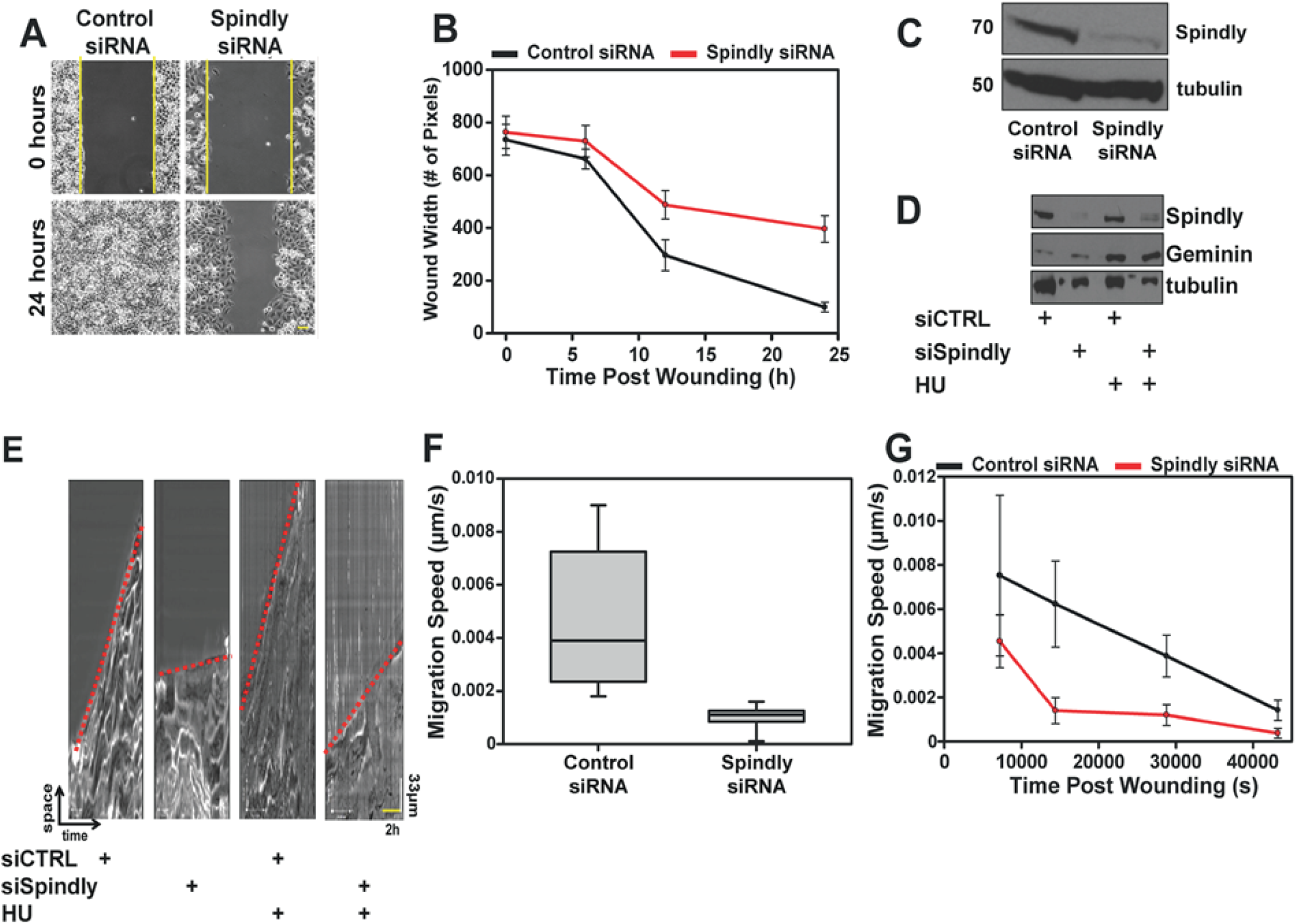
Spindly is required for rapid cell migration. A and B) U2OS cells treated with control or Spindly-specific siRNAs were plated into an ibidi silicone culture-insert inside an imaging chamber. After cells reached confluency, the insert was removed and the closure of the induced wound was followed over time using phase contrast microscopy. Three independent experiments were performed and data in the graph represent mean ± s.d. C) Western blotting confirmed that the Spindly-specific siRNAs were effective at depleting the protein. D) Spindly siRNA and control siRNA treated cells were treated with hydroxyurea (HU) to block DNA synthesis and arrest cells at the entry of S-phase. Geminin protein levels confirmed that the HU treatment worked. Tubulin used as loading control. E-G) Kymographs generated from control and Spindly-depleted cells migration movies. Red dotted lines to indicate the different slopes (i.e. velocity) between control and silenced cell. The kymographs were used to measure the speed of cells at various time points post wounding. The Spindly silenced cells were slower to migrate into a wound, regardless of whether or not HU was added to the cells. Scale bar for time and distance on the kymograph. Three independent experiments were performed and data in the graph represent mean ±s.d.

Seeing human Spindly on microtubule tips and identifying new pools of Spindly in interphase cells raised the possibility of a novel cytoskeletal role for this protein, a function different from the already established mitotic role.

### Spindly is required for cell migration

Given that we observed Spindly on microtubule tips and focal adhesions in the cytoplasm of interphase cells, and that we know that it can interact with the dynein/dynactin complex that is essential for promoting rapid cell migration, we sought to determine whether Spindly might also be involved in the cell migration process. We therefore carried out a scratch assay to study two-dimensional cell migration in Spindly-depleted cells. This method is based on the generation of an artificial gap or wound in a monolayer of cells that will drive the movement of cells on the edge of the wound to close the gap.

We silenced Spindly expression in U2OS cells by treatment with specific siRNAs for 96 hours and then seeded cells into two small wells separated by a silicon wall. Once confluence was reached, the insert was removed, generating a reproducible gap in the monolayer of cells (Figure 3A). We followed the movement of the cells on the wound edge by live imaging for at least 24 hours and then analysed the movies by measuring the width of the wound over time (Figure 3B). In multiple independent experiments, cells depleted of Spindly (typical results shown in Figure 3C) showed slow closure rates and typically were not able to close the gap after 24 hours (Figure 3B). Previous work has shown that Spindly interacts with the dynein-dynactin complex (McKenney *et al.,* 2014), and interestingly, depletion of two separate subunits of Dynactin, p150 and p50, retards wound closure migration rates (Supplemental Figure 4) comparable to Spindly-depleted cells.

Because Spindly depleted cells are defective in division and do not proliferate as well as control cells, we wished to exclude the possibility that the migration phenotype that we observed was caused by defects in cell proliferation. We therefore synchronised cells in S phase by administering Hydroxyurea (HU; 1mM) for 24 hours and repeated the scratch assay. The stabilization of geminin (Figure 3D) confirmed that the HU treatment was blocking cell cycle progression (McGarry and Kirschner, 1998). When we observed the control HU-treated and the Spindly-depleted HU-treated cells, we found that the Spindly-depleted cells HU treated were still slower to close the scratch wound.

We used kymographs to analyse the speed of cells migrating into the wound (Figure 3E) and found that the Spindly-depleted cells were slower than the control cells, regardless of whether or not cells had been treated with HU prior to wounding (Figure 3F). When we measured the velocity starting at different temporal points after wounding, we confirmed that the Spindly-depleted cells were moving slower than the controls at each time point.

To demonstrate that this phenotype was strictly due to a lack of Spindly expression, we generated a stable cell line expressing a siRNA-resistant, tet-inducible GFP-Spindly to allow the re-expression of Spindly after siRNA depletion of the endogenous protein. Figure 4 A-C shows that re-expressing an exogenous copy of Spindly rescued the migration phenotype; cells expressing wild-type GFP-Spindly showed a rate of wound closure similar to control cells, even though the overall amount of GFP-Spindly was lower than the combined level of endogenous and GFP-Spindly expressed in control siRNA treated cells. This experiment provides further evidence that Spindly depleted cells are intrinsically defective in cell migration, and that the defects we measure are not produced by off-target effects of the siRNAs used.

**Figure 4.**
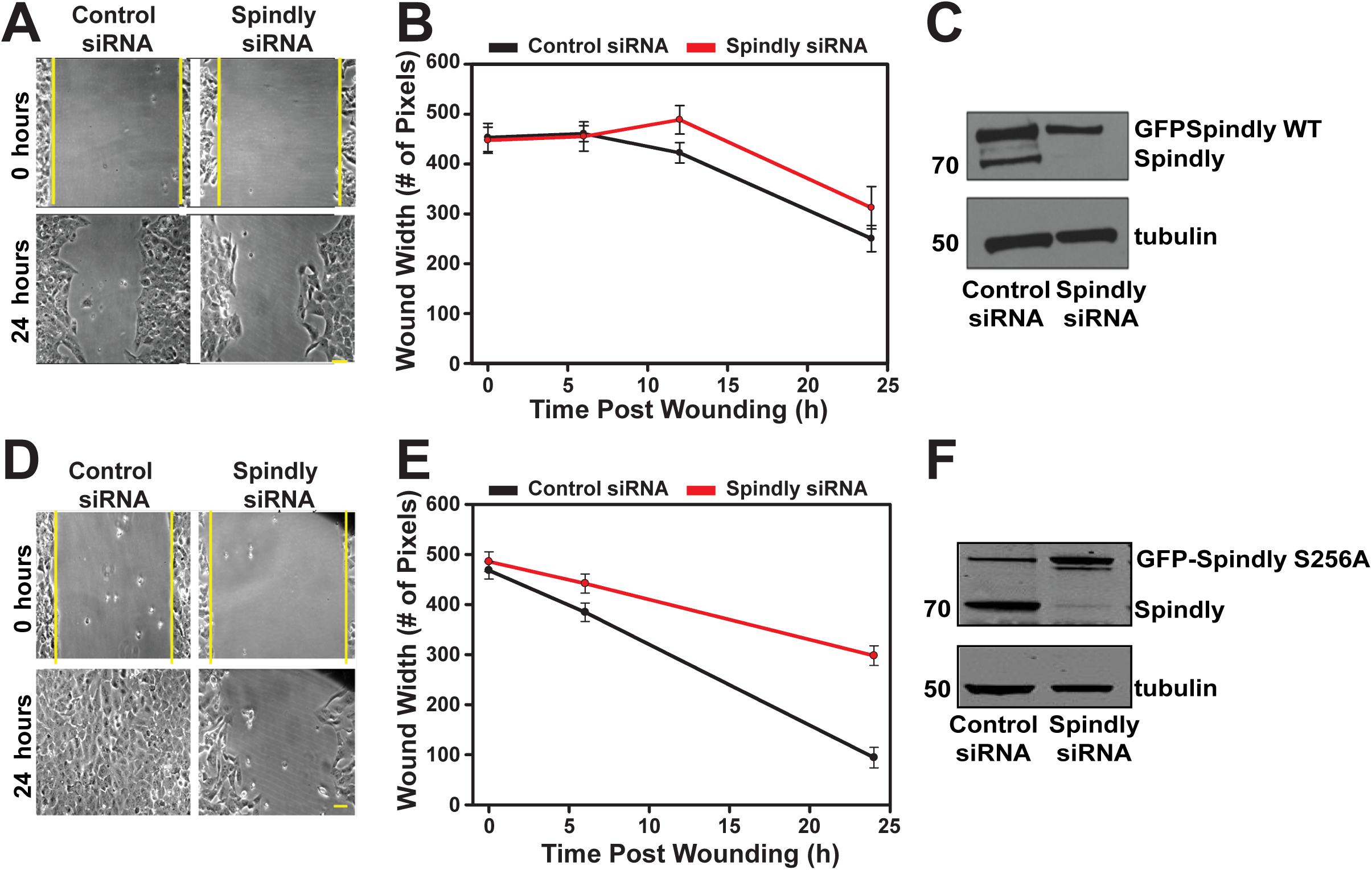
Spindly requires an interaction with dynactin to promote cell migration. U2OS cells expressing GFP-Spindly either WT or mutant (S256A) upon administration of doxycycline, were treated with control siRNA or siRNA that targets the endogenous Spindly. A-C) The expression of close to endogenous levels of GFP-Spindly WT was sufficient to nearly completely rescue wound closure. D-F) Conversely the expression of GFP-Spindly S256A did not recover the migration velocity rate.

We next wanted to determine if the non-mitotic Spindly function could be due to its association with the dynein/dynactin motor complex. To test this, we generated a stable cell line expressing a siRNA-resistant GFP-Spindly where serine 256 is mutated to alanine, a mutation that abolishes the ability of the protein to bind to dynactin (as previously described, Gassmann *et al*., 2010). We depleted the endogenous form of the protein and repeated the rescue experiment with cells that now inducibly express GFP-Spindly S256A. The expression of this mutant in the absence of the endogenous protein did not rescue the wound closure rate to control levels (Figure 4 D-F). From this data we hypothesize that Spindly is playing a role in cell migration via regulation of the dynein/dynactin motor complex.

### Spindly depletion does not grossly affect the distribution of myosin or actin filaments or centrosomal reorientation in migrating cells

There was the possibility that the depletion of Spindly from migrating U2OS cells could dramatically alter the actin-myosin or microtubule cytoskeleton as it did in Drosophila S2 cells, leading to the slow migration phenotype (Griffis et al., 2007). To check this, we fixed and stained migrating control siRNA-treated and Spindly-depleted cells with fluorescent phalloidin and an antibody that recognizes the phosphorylated (active) form of the myosin regulatory light chain. We did not observe any gross defects in the organization of actin filaments and active myosin in these cells, although there was a trend towards lower levels of phospho-myosin in Spindly-depleted cells (Figure 5A).

**Figure 5.**
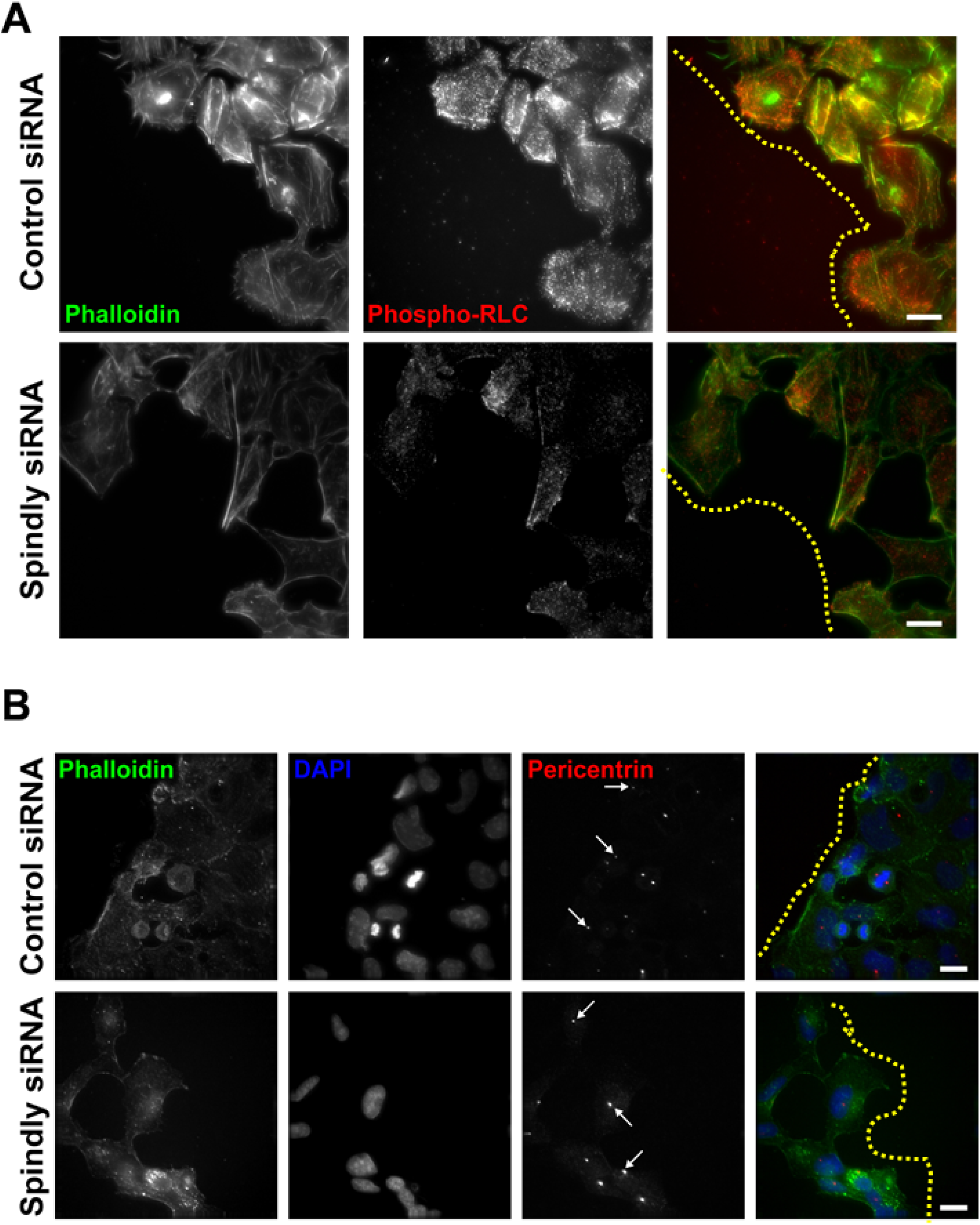
Cells lacking Spindly do not show dramatic alterations in their actomyosin cytoskeleton or in their ability to reorient their centrosomes as they migrate. A) Confluent U2OS cells that had been treated with the given siRNAs were wounded and then fixed and stained to visualize actin filaments and the active form of the RLC. Yellow dotted lines are to limit the leading edge of the cell sheet. B) Cells treated in a similar manner as in A were fixed and stained to visualize filamentous actin, nuclei, and centrosomes. Arrows show cells where the centrosome has been reoriented and is lying between the nucleus and the leading edge. Scale bars = 20μm.

An important step in polarizing migrating cells is the reorientation of centrosomes towards the leading edge, which is a dynein/dynactin dependent process and occurs after wounding (Schmoranzer et al., 2009). We wished to determine if this process was compromised in cells lacking Spindly, and so we fixed and stained control and Spindly-depleted cells four hours after scratch-wounding to visualize centrosomes and nuclei. We saw many examples of Spindly-depleted cells at the wound edge where the centrosomes were oriented correctly between leading edges and nuclei (Figure 5B). We therefore conclude that the migration defects that we observed in the Spindly depleted cells are dynein/dynactin dependent, but they are not caused by a wholesale loss of dynein/dynactin function. This is in keeping with previous results, where we found that the depletion of Spindly did not cause the redistribution of Rab5 positive early endosomes in interphase S2 cells that was seen when dynein was depleted (Griffis et al., 2007).

Spindly first emerged as a protein in human, *Drosophila* and *C. elegans* that is required for the recruitment of dynein/dynactin to kinetochores (Griffis et al., 2007, Gassmann et al., 2008, Yamamoto et al., 2008). However, other functions for Spindly appeared to have diverged. In human cells and *C. elegans*, the depletion of Spindly led to severe chromosome alignment phenotypes, but did not inhibit the shedding of spindle assembly checkpoint (SAC) proteins from aligned kinetochores (Griffis et al., 2007, Gassmann et al., 2008). In contrast, the depletion of Spindly from *Drosophila* cells did not markedly inhibit chromosome alignment, but did lead to the accumulation of SAC proteins on aligned kinetochores (Griffis et al., 2007). How much of these differences are due to species specific changes in the mechanisms of SAC silencing or the roles of dynein/dynactin has yet to be determined. Recent work showed that dynein/dynactin complex formation and processivity is facilitated by accessory factors like Spindly, which control the localization, activation and/or cargo-binding of the complex (Schroeder and Vale, 2016, Schlager et al., 2014, McKenney et al., 2014).

Just as the role of Spindly in mitotic cells appeared to be different among different species there is also an interesting bifurcation in Spindly’s interphase function. In human cells, the protein appeared to be almost exclusively nuclear, while in *Drosophila* cells, Spindly bound to microtubule plus-ends and its depletion produced a striking morphological phenotype (Griffis et al., 2007). *Drosophila* Spindly is 175 amino acids longer than the human protein and contains four clusters of positively charged residues (consensus sequence: TPAKPQ-L/R/M-KGTPVK) that could potentially interact with the negatively charged C-terminal tubulin tails. Interestingly, embedded in these four C-terminal repeats are seven consensus CDK1 phosphorylation sites; modification of these sites could reverse these charge-charge interactions and explain why this protein is not seen on microtubules in mitotic cells. A recent paper showed that altering the levels of Spindly in migrating border cells in the oocyte alters their migration speed (Clemente *et al*., 2017), suggesting that a role in migration is shared between *Drosophila* and human Spindly.

Human Spindly is primarily a nuclear protein in interphase cells, and before this report had not been shown to have any interphase role. Human Spindly lacks the charged regions seen in the *Drosophila* protein and is much less regularly observed on microtubules. Therefore, the particles that we observed moving on plus-ends or near the cortex were probably not directly bound to microtubules, but could be associated either with vesicles, plus-tip proteins, or other microtubule binding complexes. A recent publication has reported that CENP-F can also be seen associating with microtubule plus-ends, and so it may be that a subset of kinetochore proteins associate with microtubule cargoes to help them attach to plus-ends (Kanfer et al., 2017). Additionally, Spindly can now be considered like Zw10 and Mad1 as a kinetochore proteins that has additional roles outside of mitosis (Wan et al., 2014, Schmitt, 2010).

Given our data that the wild-type Spindly protein can rescue migration but the S256A dynactin-binding deficient point mutant cannot, we favour a model in which one or a few particular dynein/dynactin functions in migrating cells require Spindly. Observing Spindly enriching in the proximity of focal adhesions at the leading edge, may also indicate a role for Spindly in cell adhesion, a process that has already been shown to be influenced by dynein/dynactin activity (Rosse et al., 2012). Future experiments to measure cell adhesion and/or contractility in the absence of Spindly will be needed to test whether Spindly has a role in regulating this process. However, this report represents a significant advance in our understanding of Spindly’s role in interphase cells.

### Materials and Methods

#### Cell culture, transfection and RNA interference

U2OS and human foreskin fibroblast (FB) cells (a gift from Prof. A. Huebner, Technische Universitat Dresden, Dresden, Germany) were maintained in Opti-MEM (Thermo Fisher Scientific; Waltham, MA, USA) supplemented with 10% heat-inactivated fetal bovine serum (HI-FBS) (Thermo Fisher Scientific), 1% Penicillin, Streptomycin and Glutamine (Thermo Fisher Scientific) for no more than 30 passages. Cell lines were grown at 37°C with 5% CO_2_ in a humidified incubator.

To generate stable cell lines expressing wild-type or mutant (S256A) GFP-Spindly, U2OS cells with an integrated FRT site and a Tet repressor (a gift from Prof. A. Lamond, University of Dundee), were co-transfected with the pOG44 vector (a gift from Prof. J. R. Swedlow, University of Dundee) together with the pCDNA5/FRT/TO LAP-Spindly constructs (Gassmann et al., 2010) in a ratio of 9:1 pOG44:pgLAP vector. Stable integrating cell lines were subsequently drug selected in media containing 150 μg/mL of Hygromycin (MilliporeSigma; Billerica, MA, USA) and 15 μg/mL of Blasticidin, and clonally isolated. Expression of the GFP construct was induced by administration of doxycycline (0.1-μg/mL; MilliporeSigma).

DNA transfection procedure was carried out using the FuGene HD reagent (Promega; Madison, WI, USA) according to manufacturer’s instructions. Cells were transfected 24 hours after being seeded with a FuGene/DNA ratio of 3:1 incubated in 200μL of serum-free media for 30 minutes at room temperature. The mix was dropped onto cells growing in OptiMEM supplemented with 10% FBS and plates were incubated at 37°C for at least 24 hours before experiments were conducted. All the DNA plasmids used in this study were purified using the QIAprep Spin Miniprep Kit (QIAGEN; Hilden, Germany), following manufacturer’s instructions.

Small interfering RNA oligonucleotides were synthesized by SIGMA and transfected into cells using Lipofectamine RNAiMax (Thermo Fisher Scientific) according to the manufacturer’s instructions. The oligonucleotide sequences used for siRNA knockdown are as follows: a GC-matched non-targeting control (MISSION Negative Control, SIGMA; SigmaMillipore) diluted to a final concentration of 20nM; Spindly Endo1 (GAAAGGGUCUCAAACAGAA) and Spindly-UTR-66 (CUUGAUCUGACAUAUAUCA) (neither of which target the expressed Spindly constructs) combined together to a final concentration of 20nM. Cells were seeded and directly treated. Treatment was left on for 96 hours and then cells were either fixed, harvested or seeded again for the subsequent analysis.

#### Western blotting and cell fractionation experiment

To perform the immunoblot analysis, cells were lysed using the following lysis buffer: 50mM Tris/HCl pH 7.5, 150 mM NaCl, 100mM N-Ethylmaleimide (NEM), 0.3% CHAPS, 1mM EGTA, 1mM EDTA, 10mM Na-β-glycerophosphate, 1mM Na-orthovanadate, 50mM Na-Fluoride, 10mM Na-pyrophosphate, 270mM sucrose, 0.1mM PMSF, 1mM Benzamidine, 0.1% β-mercaptoethanol, 1 protease inhibitor cocktail tablet (Roche Diagnostics, Basel, Switzerland) for 10 mL of buffer. Cells were then transferred into Eppendorf tubes and put under constant agitation for 10 minutes at 4°C. Samples were subsequently spun down for 15 minutes at 13,000 rpm at 4°C. Supernatants were collected and stored at −80°C. Protein concentration was measured using Bradford dye (BioRad; Hercules, CA, USA), according to the manufacturer’s instructions. Samples were prepared in 2x loading buffer (Novex LDS sample buffer; Thermo Fisher Scientific) and boiled at 95°C for 5 minutes. Soluble fractions were resolved on Tris-glycine SDS-PAGE gels (4-12% gradient gel; Thermo Fisher Scientific) and transferred to a nitrocellulose membrane. Spindly, tubulin, geminin, GAPDH and PCNA were detected using specific antibodies: rabbit anti-Spindly (Griffis et al., 2007), mouse anti-PCNA, mouse anti-GAPDH, and rabbit anti-geminin from Santa Cruz Biotechnology (Dallas, TX, USA), rat anti-tubulin from Thermo Fisher Scientific, sheep anti-GFP from Novus Biological (Abingdon, UK). The proteins were then visualised using ECL solution Thermo Fisher.

To perform cell fractionation U2OS cells were grown to confluence in a 150mm petri dish, washed with chilled PBS 1X and lysed in 200μL of Buffer A (10 mM HEPES, pH7.9, 10 mM KCl, 1.5 mM MgCl_2_, 1 mM DTT, 0.5 μg/μL Leupeptin, 0.5 μg/μL Aprotinin, 0.5 μg/μL Pepstatin, 0.1 mM PMSF, 0.34 M sucrose, 10 % Glycerol, 0.1 % Triton X-100, 1μL DTT, 1μL LPC), swirled and placed on ice for 20 minutes. Cells were then scraped, transferred to Eppendorf tubes and spun down at 1,400 RCF for 10 minutes at 4°C. Supernatant was removed and collected into a new Eppendorf as the “cytoplasmic” fraction. To the remaining pellet we added 50μL of Buffer A combined with 0.2 μL benzonase nuclease (SigmaMillipore) and 10 μL 0.1M CaCl_2_ and incubated at 37°C for 1 minute. The sample was then placed back on ice and supplemented with 0.2μL of 0.5M EGTA and incubated for 5 minutes. We then spun down the sample at 1,400 RCF for 5 minutes at 4°C. The supernatant was removed and collected into a new Eppendorf tube as the “nuclear” fraction.

#### Immunofluorescence microscopy

Cells were seeded onto sterilised glass coverslips (Menzel-Glaser, Braunschweig, Germany). When confluent, cells were fixed in 4% PFA in 1x PHEM (60mM PIPES, 25mM HEPES, 10mM EGTA, 2mM MgCl_2_, pH 6.9) for 10 minutes at room temperature, washed in PHEM-wash (1xPHEM + 0.1% Triton X-100), permeabilized in PHEM-T (1xPHEM + 0.5% Triton X-100) for 5 minutes and fixed again in 4% PFA for 10 minutes. Blocking of non-specific antigen recognition was performed in Abdil (1xTBS-0.1% Tween, 2% BSA) for 1 hour. At this point coverslips were incubated with primary antibodies for one hour at room temperature or overnight at 4°C (antibodies were diluted in Abdil). The following antibodies and probes were used: phalloidin-Atto 488 (SigmaMillipore), mouse anti-p50 (BD-Biosciences; Oxford, UK), rabbit anti-pMyosin light chain (Cell Signaling Technology; Danvers, MA, USA), rabbit anti-pericentrin (SigmaMillipore) and rabbit anti-Spindly (Griffis et al., 2007); anti-vinculin (Sigma). Following incubation for one hour with cross-subtracted secondary antibodies (either AlexaFluor-labelled (Thermo Fisher Scientific) or Cy3 or Cy5-labelled (Jackson Immunoresearch; West Grove, PA, USA)) diluted in Abdil with DAPI (SigmaMillipore). Antibody incubations were followed by three washes with PHEM-wash. Coverslips were mounted in Dako fluorescence mounting medium (Carpinteria, CA, USA). Images were collected using a fluorescence microscope (DeltaVision Elite; GE Healthcare Life Sciences, Issaquah, WA, USA). Images were then processed using OMERO software.

#### Cell migration assay and scratch assay

U2OS cells were treated for 96 hours with either negative control or Spindly specific oligonucleotides, seeded into a Silicone Culture-Insert (ibidi; Matinsried, Germany) set into a 35mm μ-Dish (ibidi) and left to grow for at least 24 hours. Once cells were confluent, the insert was removed and cells were washed once with fresh media and then the media was replaced with a CO2 independent media (Leibovitz’s L-15, supplemented with 10% FBS and 1x Penicillin/Streptomycin/Glutamine; Thermo Fisher Scientific). A sample of the cell population was collected for Western Blotting to confirm the silencing of Spindly. For blocking the cell cycle, cells were treated with hydroxyurea (HU) (1mM; SigmaMillipore) for 24 hours before being imaged, and a sample of the cell population was collected for Western Blotting to confirm silencing of Spindly and S-phase synchronisation. For the rescue experiments U2OS cells stably and inducibly expressing WT or S256A GFP-Spindly were treated with siRNAs that targeted either the endogenous Spindly or nothing. Subsequently, doxycycline (100ng/mL) was administrated overnight to induce expression of the GFP construct, and a small portion of the cell population was harvested for Western Blotting to observe depletion of the endogenous Spindly and the expression of the GFP-tagged proteins.

Imaging was performed on a Nikon Ti microscope (Nikon, Tokyo, Japan) fitted with an environmental control chamber (Okolab, Pozzuoli, Italy), 20x 0.45NA objective, Nikon PerfectFocus System, and a Photometrics Cascade II camera (Photometrics, Tucson, AZ, USA) and NIS Elements software. Images were then processed using Fiji software. Measurements were conducted by drawing a line between the edges and measuring the distance at the indicated time points. Kymograph analyses were conducted using Volocity software (Perkin Elmer; Waltham, MA, USA) to measure the velocity of movement over the time.

Human fibroblast cells were seeded onto glass coverslip and left grow until confluent. Then a scratch was mechanically generated by using a 200μL tip and slides were fixed at different times to check for protein recruitment at the leading edge. Staining was performed as above.

#### TIRF Imaging

Cells were grown and transfected in ibidi μ-Dishes; two hours prior to imaging, the media was removed and replaced with a CO_2_ independent Leibovitz’s L-15 media without phenol red to lower autofluorescence. Imaging was performed on a Nikon Ti-E microscope with an environmental control chamber (Okolab), a PAU/TIRF slider, 63x and 100x 1.49 N.A. objectives, PerfectFocus system, a custom-built four-color (405nm, 488nm, 561nm, 647nm) diode laser (Coherent Inc., Santa Clara, CA, USA) system that has a Gooch and Housego (Ilminster, UK) AOTF shutter (Solamere Technology; Salt Lake City, UT, USA), an emission filter wheel (Nikon) with appropriate filters for eliminating crosstalk between channels (Chroma Technology Corp, Bellow Falls, VT, USA) and a Photometrics Evolve Delta camera (Tucson, AZ, USA). Images were all captured with μ-Manager (Open Imaging Inc., San Francisco, CA, USA). In order to better visualize the moving particles of Spindly among the basal fluorescence, we utilized a method that was developed for tracking particles within Drosophila oocytes (Parton et al., 2011). Briefly, images were denoised with a 3D Gaussian blur filter with a radius of 1 pixel. A temporal median filter was then used to extract only moving “foreground” features using a sliding time window (half-width = 4), and “foreground” set to 4 standard deviations over the median value. The outputs of the temporal filter were then trailed using a sliding window of time-points (half-width = 2) and averaged to make moving particle trails obvious in all still frames. The temporal median filter plugin and trails plugin were created by Graeme Ball (https://github.com/graemeball/IJ_Temporal).

## Acknowledgments

We dedicate this paper to the memory of Dr. Michael Davidson and to all of the contributions that he made to the field of cellular imaging. We thank the Light Microscopy Facility, College of Life Sciences, University of Dundee, and especially Alan Prescott and Graeme Ball for help with imaging and image analysis. We thank Yu-li Wang for the gift of the RFP-Zyxin construct and Jason Swedlow, Angus Lamond, and Angela Huebner for providing other cell lines and contructs. We thank Julian Blow for providing reagents and advice for the hydroxyurea arrest assay. C.C. was supported by a non-clinical studentship from Cancer Research UK within the Dundee Cancer Centre (grant number: C5314/A11784). ERG was supported by a Wellcome Trust RCDF award (090064/Z/09/Z), and a Wellcome Trust Strategic award to the Centre for Gene Regulation and Expression (097945/B/11/Z).

## Supplementary Information

**Figure S1: Spindly is not an exclusively nuclear protein**. U2OS cell lysate fractionation was performed and fractions were probed with multiple antibodies to confirm the proper isolation of the different fractions. The blot confirms that Spindly is not seen in the chromatin fraction. In the nuclear fraction (middle) we observe PCNA, and Spindly enrichment along with Histone H3. Spindly is also found in the cytoplasmic fraction, which is confirmed by the presence of GAPDH.

**Figure S2: Spindly localization at the leading edge of migrating cells requires actin filaments**. Human fibroblasts were grown to confluence and then a scratch-wound was made in the monolayer. Cells were allowed to migrate for two hours before they were treated with either Latrunculin A (LAT A 100 nM) or Jasplakinolide (JAS 100nM) and then fixed 20 minutes later and stained to visualize Spindly, p50, and actin. Scale bar = 20μm

**Figure S3: Spindly co-localizes with Zyxin at focal adhesions**. A confluent monolayer of U2OS cells transfected with RFP-Zyxin were wounded, fixed, and stained to visualize Spindly and focal adhesions. Co-localization of Spindly with Zyxin was observed at the basal cell cortex, but only at peripheral focal adhesions. Arrows show regions where Spindly and Zyxin colocalize.

**Figure S4: Depletion of dynactin inhibits cell migration**. A) U2OS cells treated with control or Dynactin- specific siRNAs (either p150 or p50) were plated into an ibidi silicone culture-insert inside an imaging chamber. After cells reached confluency, the insert was removed and the closure of the induced wound was followed over time using phase contrast microscopy. B) Quantification of the width of the scratch in control and dynactin depleted U2OS cells over time. Data indicate the mean ± s.d. from at least 3 independent experiments. C) Western blotting of the same population of cells confirms the silencing; Tubulin (or Actin) was used as loading control.

**Movie S1: Spindly can be observed moving along tracks on the cell cortex**. U2OS cells stably expressing GFP-Spindly were imaged using total internal reflection (TIRF) microscopy. The movie on the left shows the original data. To better visualize moving particles, a temporal median filter was used to highlight particles that moved from frame to frame and a trailing function was used to link particles. The processed image is shown on the right. Movie is shown at 20x realtime. Field of view = 40.72 × 40.72 μm.

**Movie S2: Spindly is seen at the basal cortex in cells moving into a wound**. U2OS cells stably expressing low levels of GFP-Spindly that were at the end of a scratch-wound were observed in TIRF as they migrated into the wound. Movie is shown at 600x realtime. Field of view = 131.2×131.2 μm.

**Movie S3: Spindly can be observed colocalizing with dynamic microtubule tips**. U2OS cells stably expressing GFP-Spindly (Green) were transfected with TagBFP2-Tubulin (Red) and imaged in TIRF. Movie is shown at 50x realtime. Field of view = 30.1 × 30.μm.

**Movie S4: Spindly stably associates with the leading edge and to regions proximal to focal adhesions in migrating cells**. U2OS stably expressing GFP-Spindly and transiently transfected with RFP-Zyxin were imaged in TIRF. Movie is shown at 600x realtime; field of view = 31.4 × 31.4 μm.

